# CRMnet: a deep learning model for predicting gene expression from large regulatory sequence datasets

**DOI:** 10.1101/2022.12.02.518786

**Authors:** Ke Ding, Gunjan Dixit, Brian J. Parker, Jiayu Wen

## Abstract

Recent large datasets measuring the gene expression of millions of possible gene promoter sequences provide a resource to design and train optimised deep neural network architectures to predict expression from sequences. High predictive performance due to the modelling of dependencies within and between regulatory sequences is an enabler for biological discoveries in gene regulation through model interpretation techniques.

To understand the regulatory code that delineates gene expression, we have designed a novel deep-learning model (CRMnet) to predict gene expression in *Saccharomyces cerevisiae*. Our model outperforms the current benchmark models and achieves a Pearson correlation coefficient of 0.971. Interpretation of informative genomic regions determined from model saliency maps, and overlapping the saliency maps with known yeast motifs, support that our model can successfully locate the binding sites of transcription factors that actively modulate gene expression. We compare our model’s training times on a large compute cluster with GPUs and Google TPUs to indicate practical training times on similar datasets.

## 1 INTRODUCTION

Cis-regulatory sequences also referred to as cis-regulatory modules (CRMs), are composed of promoters, enhancers, silencers, and insulators (Davidson and Erwin, 2006). The DNA-binding regulatory proteins, transcription factors (TF), identify and bind to particular cis-regulatory sequences to control gene expression (Ni and Su, 2021). Alterations to the cis-regulatory sequences will influence the interaction with transcription factors, thereby influencing cell phenotype and cell-state transitions (de Boer et al., 2020). Increasing evidence demonstrates the significance of cis-regulatory element modification in relation to numerous diseases, such as cancer and diabetes (Mathelier et al., 2015). Thus, understanding how cisregulatory elements regulate gene expression has become critical for us to understand transcriptional gene regulation. However, it has been extremely difficult to directly predict the expression of DNA sequences due to a lack of high quality data. Recently, more than 100 million random promoter sequences and their corresponding expression levels have been identified in a high-throughput manner by measuring the expression output of the sequences regulating yeast gene constructs using Gigantic Parallel Reporter Assay (GPRA) (de Boer et al., 2020). This dataset provides the large training sets necessary for developing models to help decode the cis-regulatory logic. Consequently, sequence-to-expression models have been proposed to predict how changes in cis-regulatory sequences will affect gene expression (Vaishnav et al., 2022).

This explosion of genomics data size presents a challenge to conventional analysis methods, and the subtle long-range interactions in genomic data challenge the explicit feature engineering stage required in other predictive modelling approaches. Deep learning, utilising deep neural network (DNN) models, which benefits greatly from large datasets, has therefore increasingly found application in genomics. Composed of multiple (“deep”) processing layers such as convolutional, recurrent, and dense layers, deep learning models can learn complex patterns and features in large datasets at various abstraction levels by combining such processing layers into appropriate DNN architectures.

Concurrently with the research and development of increasing deep and complex DNN architectures, the development of parallel acceleration hardware such as graphical processing units (GPUs) and tensor processing units (TPUs), and high performance clusters/cloud computing, enables the training of these more complex models by reducing the overall training time (Wang et al., 2020). As neural networks are becoming more sophisticated and the volume of scientific data keeps growing, the model training time on these different high performance computing (HPC) architectures is an important issue.

In this study, we propose a novel DNN model (CRMnet), a Transformer encoded U-Net, for predicting the expression levels of yeast promoter DNA sequences, which achieves a Pearson correlation coefficient of 0.971 in the test dataset, improving upon the benchmark models proposed in (Vaishnav et al., 2022). By accurately predicting the expression from promoter sequences, such models can be used predictively to design new regulatory sequences in synthetic biology, study the predicted effects of mutations (Vaishnav et al., 2022), and, by interpreting the model, help in understanding the determinants of gene regulation. Here we interpret the model by visualizing saliency maps, showing we are able to identify key regions in the promoter sequences which most affect the corresponding expression. We demonstrate that our model can learn biologically meaningful information by quantifying the saliency information over known yeast sequence motifs. We compare the performance of our model on large datasets on parallel hardware of graphical processing units (GPUs) and tensor processing units (TPUs) on a HPC cluster.

## 2 CRMNET: SEQUENCE-TO-EXPRESSION DEEP LEARNING MODEL

The study of (Vaishnav et al., 2022) experimentally determined the gene expression driven by millions of random promoter sequences. This was performed by embedding random 80-bp DNA sequences in a promoter construct with the resultant expression assayed in yeast (S. cerevisiae) using high-throughput sequencing.

In this study, we propose an improved novel DNN model to predict the measured expression level of yeast promoter DNA sequences with higher performance than the convolutional neural network (CNN) and transformer deep neural network models proposed in (Vaishnav et al., 2022). In this DNN architecture, we propose a transformer-encoded U-Net (CRMnet). Analogous to the original U-Net model (Ronneberger et al., 2015), our deep learning model has an initial encoding stage that extracts feature maps at progressively lower dimensions, optimised for the detection of features such as transcription factor binding sites (average length of approximately 11bp), and a decoder stage that upscales these feature maps back to the original sequence dimension, whilst concatenating with the higher resolution feature maps of the encoder at each level to retain prior information despite the sparse upscaling. This approach decodes a feature map at base-level precision. Recent work on transformer architectures has shown that their attention mechanisms can extract more global dependency information compared with convolutional layers (Dosovitskiy et al., 2020); therefore, we included a transformer encoder stage after the convolutional layer to extract this information.

Our model thus consists of four components (Figure 1): 1D convolutional neural network-based encoders to extract neighboring features in the input DNA sequences, a transformer encoder to extract longer range dependencies in the input sequence, 1D convolutional neural network-based decoders, with skip connections, to project the extracted features to the original sequence input dimension, and a multi-layer perceptron to predict the expression levels from the extracted features.

**Figure 1:**
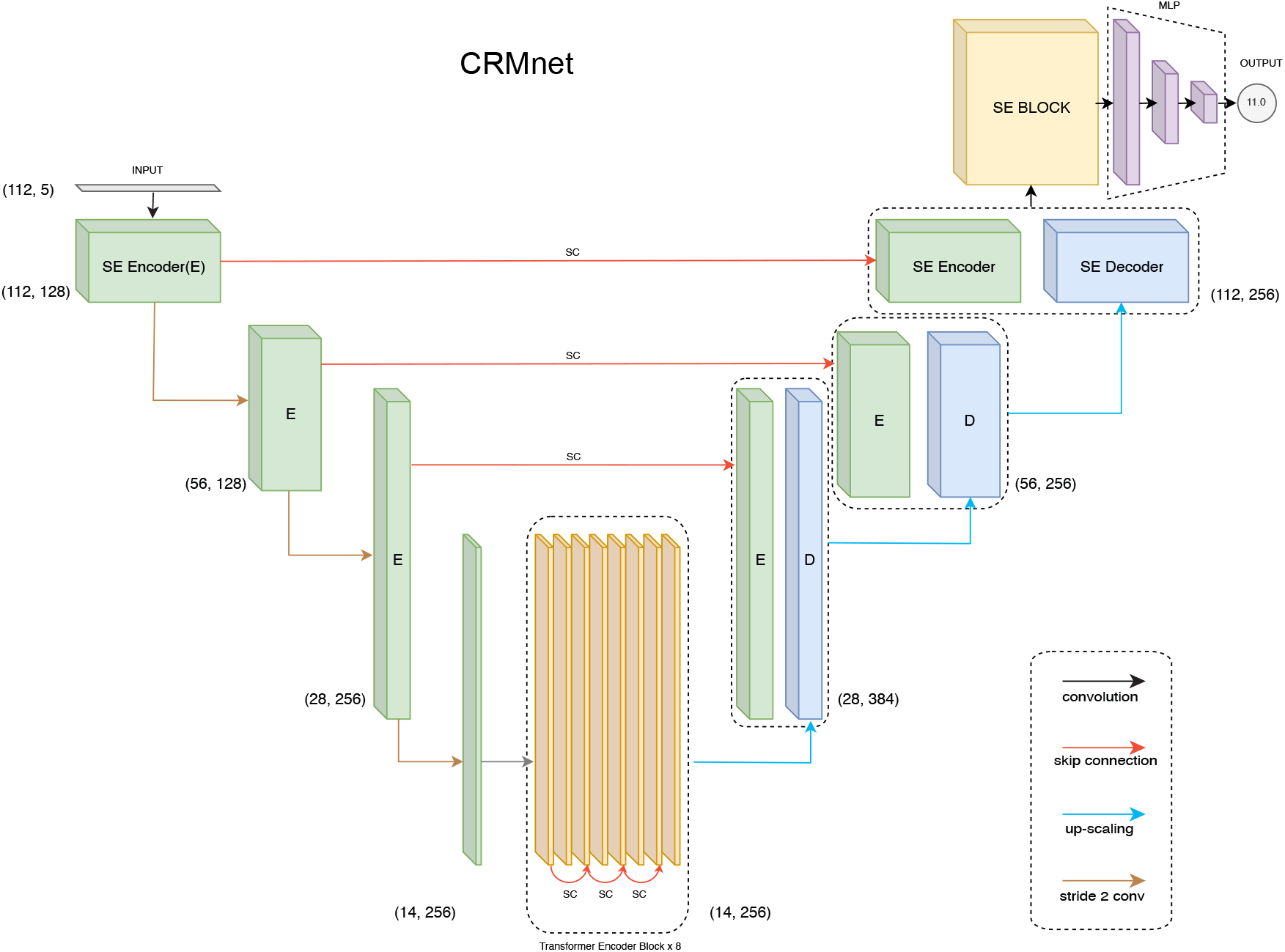
CRMnet model’s architecture. Our CRMnet consists of Squeeze and Excitation (SE) Encoder Blocks, Transformer Encoder Blocks, SE Decoder Blocks, SE Block and Multi-Layer Perceptron (MLP). Similar to the UNet architecture, the encoder and corresponding decoder at the same level have a skip connection (SC) so the decoder utilizes the concatenation of the upsampled feature map with the corresponding higher resolution encoder feature map at that level.

### 2.1 1D CNN-based Encoder

Our CNN-based encoder is built to extract features from genomic data inputs. The input is onedimensional length 112bp DNA sequences (80bp promoter sequence plus padding, see Methods) which is one-hot encoded (A,C,G,T + N for padding). 1D convolutional layers then learn filter parameters to extract predictive features from combinations of adjacent bases, with a filter size set to 11 in order to cover the average length of transcription factor motifs (Stewart et al., 2012). Additionally, we add a squeeze excitation layer after each 1D convolutional layer because the squeeze and excitation operation has been demonstrated to improve the overall performance of CNN-based models by assigning importance scores to the different feature maps (Hu et al., 2018). Moreover, the original U-Net encoder block (Ronneberger et al., 2015) is modified by performing the down-sampling with a stride two convolution operation instead of max pooling, as the additional model parameterisation has been shown to improve performance (Springenberg et al., 2014).

### 2.2 Transformer Encoder

The transformer encoder accepts the CNN’s down-scaled feature maps as input. Each individual transformer encoder block follows the vanilla transformer architecture which is made up of a position-wise feed-forward network and a multi-head self-attention feed-forward network (FFN) module (Vaswani et al., 2017). Within each module, residual/skip connections and layer normalization are utilized in order to train a deeper neural network.

Unlike convolutional neural network stages which implicitly extract dependency information in local neighbourhoods through the use of fixed-size kernels, transformer encoders use self-attention to extract global dependency information across the inputs, while explicitly encoding the positional information embedded into the input. Similar to other transformer-encoded models, we embed the positional information of the down-scaled input vectors (length 14) to our transformer encoder (Chen et al., 2021). We represent the positional information using sinusoidal position encodings and add it to the input token before feeding it to the transformer encoder.

### 2.3 1D CNN-based Decoder

The CNN-based decoder blocks are very similar to the original U-Net decoders, which use up-sampling to learn the representation (Ronneberger et al., 2015). Using a 1D transpose convolution operation to upsample the resolution and attaching a squeeze excitation layer after each convolutional layer to up-weight the critical feature maps are the main differences in our implementation. The outputs are then concatenated with the skip connections from corresponding encoder levels to compensate for the potential loss of spatial information during downsampling.

### 2.4 SE Block

It has been demonstrated that using Squeeze-and-Excitation (SE) blocks can significantly improve the generalization power of CNN-based models and achieve significant performance enhancements in several state-of-the-art CNN models with negligible increase in computational cost (Hu et al., 2018). The SE Block will initially compress the input feature map generated by learned convolutional filters using global max pooling. The channel-specific statistics will then be forwarded to the excitation operation, which will utilize two non-linear, fully connected layers to highlight the key channels. In other words, the SE Block can be regarded as a channel-specific self-attention function that compensates for the inability of the convolution operator to model the relationship among channels. As a result, we decided to adopt the SE operation after each convolutional layer and added a SE block in order to empower our model to focus on channel-specific feature responses of the convolution layers.

### 2.5 MLP

Our model will learn the expression levels from the extracted features utilizing a multi-layer perceptron (MLP). The fully connected dense layer will learn the non-linear combination of the extracted features from preceding layers. In the hidden layer of MLP, we use ReLU as the activation function (where alpha equals 0.1). Then, a linear activation is used to make the prediction of the expression levels in the final output neurons. To avoid overfitting, each dense layer is followed by a dropout layer. The first two dropout layers’ dropout values are equal to 0.2 and the rest have dropout values equal to 0.1.

### 2.6 Pre-training and Fine-tuning of models

We utilized a transfer learning approach to improve the performance of CRMnet. Specifically, to utilize the largest possible training set we pre-trained a more general model on a large dataset of randomly sampled data combining datasets from yeast grown in two different media types (“complex” and “defined” from (Vaishnav et al., 2022)). We then conducted a fine-tuning training stage in which the pre-trained model is retrained on the complex medium samples only, as used in the test sets of our study. The pre-trained model’s parameters were all unfrozen and trainable. The pre-trained model weights serve as good initializations for the fine-tuning of particular datasets to improve the model’s performance on a target task (You et al., 2021). As demonstrated in Figure 4, this method of transfer learning can improve model performance.

## 3 RESULTS AND DISCUSSION

We first evaluate the predictive performance of our model and compare the performance to that of existing deep learning models. We then use ablation studies to understand the roles of the subparts of our model. To demonstrate the biological significance of our model, we further apply saliency maps for model interpretation and compare with enriched transcription factor binding site motifs discovered by probabilistic motif discovery. Finally, we compare the training time between TPUs and GPUs.

### 3.1 Performance evaluation

We here first present the predictive performance of our fine-tuned deep learning model on independent experimental test sets of both random and native (i.e. wild-type sequences found in yeast) promoter sequences in both complex and defined mediums (see Methods). To evaluate the performance of the deep learning models on the test datasets, we measured the Pearson Correlation Coefficient (*r*) and Coefficient of Determination (*R*^2^). The results show that our fine-tuned CRMnet model achieved excellent prediction performance on both native and random sequences (*r* = 0.971, and *r* = 0.987, respectively) in complex medium (Figure 2) and in defined medium (*r* = 0.955, and *r* = 0.973, respectively, Supplementary figure S1).

**Figure 2:**
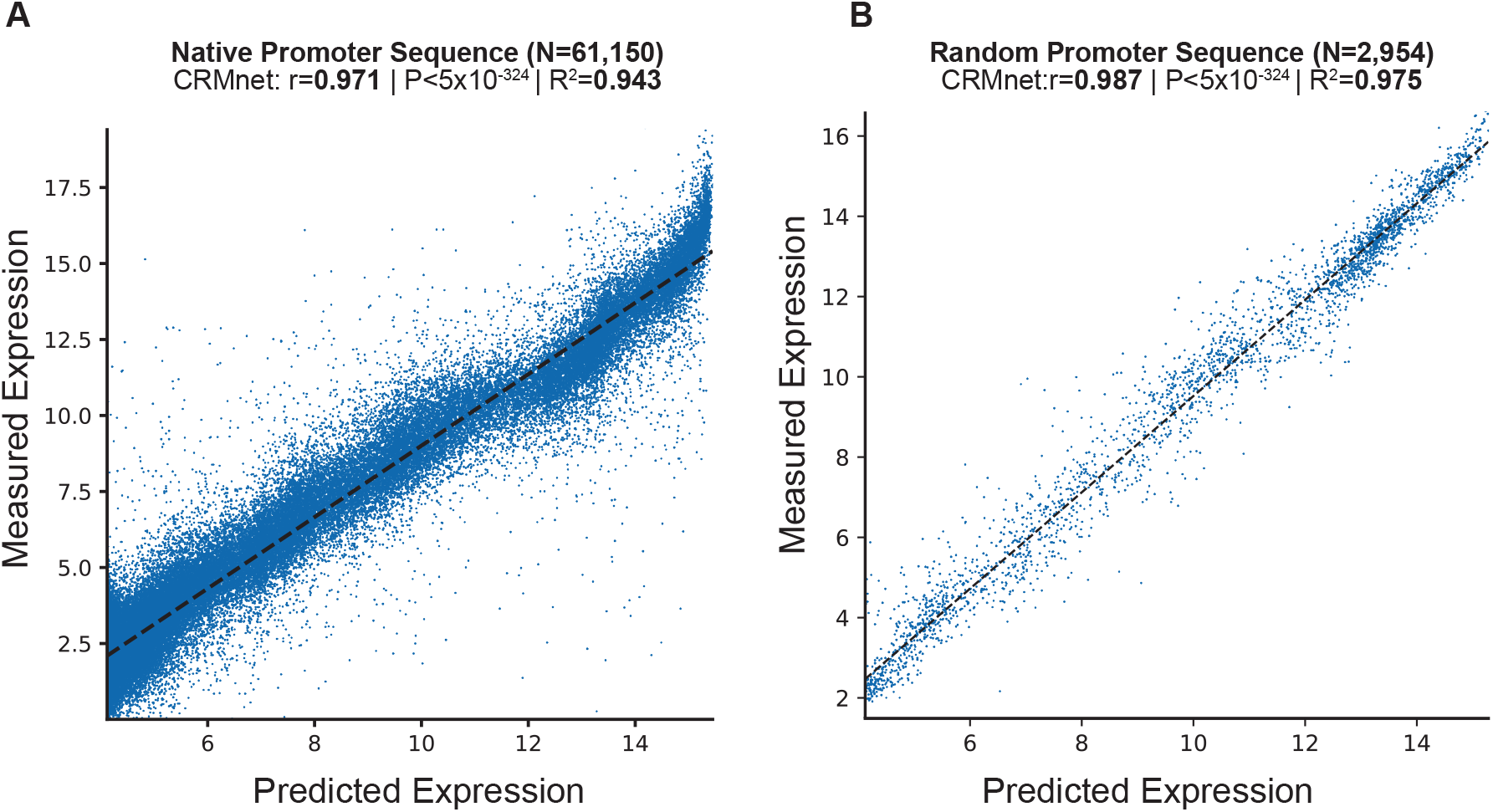
Prediction of expression from yeast native sequences from CRMnet. CRMnet tested on **A:** native promoter sequences; and **B:** random promoter sequences. The y-axes represent measured expression levels, while the x-axes represent predicted expression levels. As a benchmark, the model performance metrics of the Pearson *r* value, associated two-tailed p-values, and R-square for the transformer model from (Vaishnav et al., 2022) showed: A: r=0.963, P < 5 × 10^−324^, *R*^2^=0.927; B: r=0.978, P < 5 × 10^−324^, *R*^2^=0.95.

We next compared our model’s performance with the benchmark models on the same test datasets. Our CRMnet model outperforms the benchmark transformer model proposed by (Vaishnav et al., 2022) in both native and random promoter test datasets in both mediums (Figure 3), and also outperforms other existing deep learning models referred to in (Vaishnav et al., 2022).

**Figure 3:**
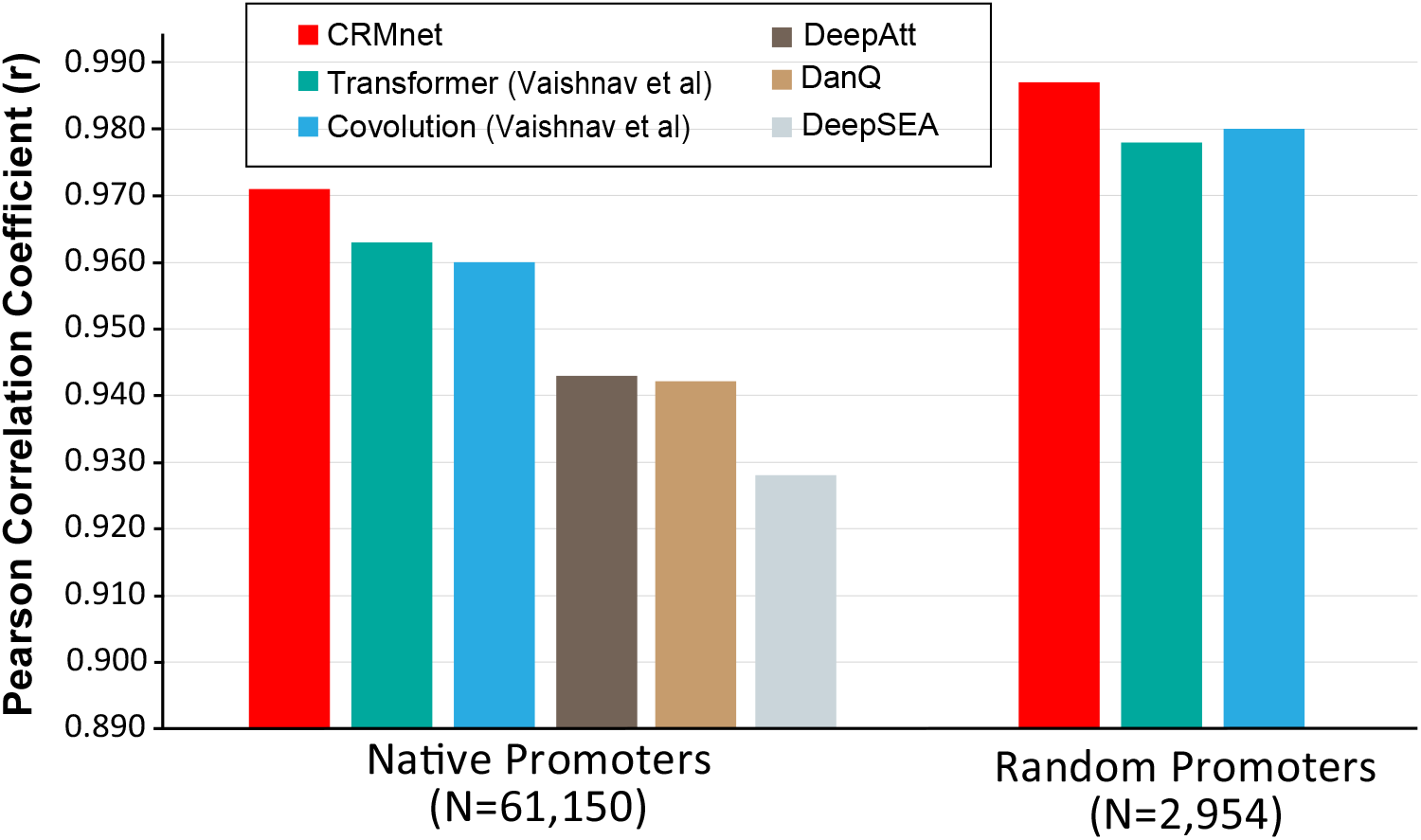
Benchmarking the CRMnet’s performance against existing neural network architectures. The prediction performance of CRMnet on yeast native promoters and random promoters was compared with the transformer and CNN models from (Vaishnav et al., 2022) and other existing DNNs (DeepAtt (Li et al., 2021), DanQ (Quang and Xie, 2016) and DeepSEA (Zhou and Troyanskaya, 2015)). The performance of DeepAtt, DanQ and DeepSEA on random promoters not published in (Vaishnav et al., 2022)

### 3.2 Ablation study

To determine the contribution to the performance of the various components of our model, we performed an ablation study. Specifically, for each ablation experiment, we constructed a new model with the ablated block removed from the original CRMnet architecture and trained the new model using the same training dataset as the original CRMnet. We then evaluated the performance of our models using the native and random test datasets (complex medium) (Figure 4).

**Figure 4:**
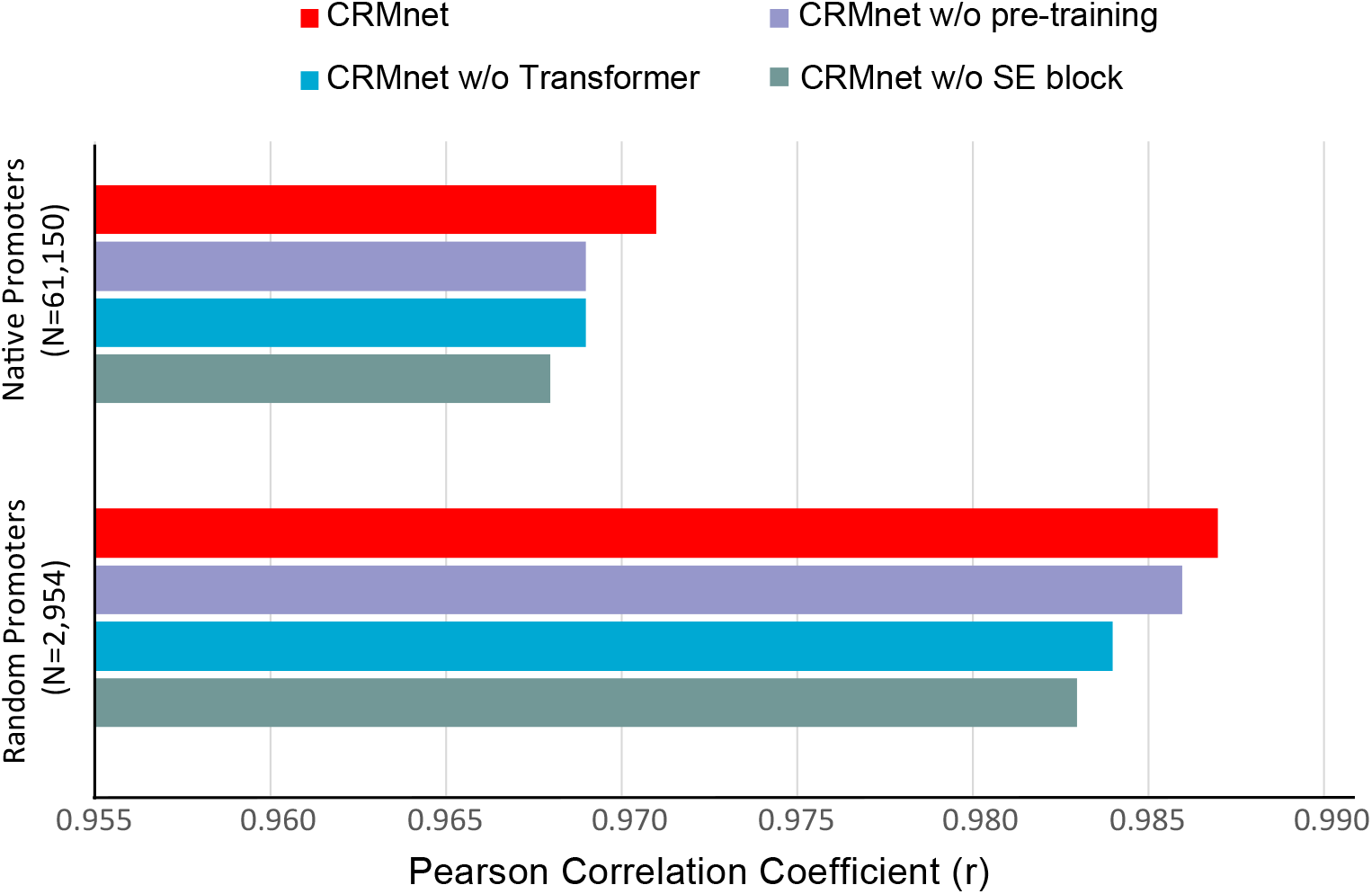
Prediction performance for ablation study. Comparisons of prediction performance of models testing on native promoters and random promoters are shown: the full model (fine-tuned CRMnet), the model without transfer learning (CRMnet without pre-training), the model without transformer block (CRMnet without transformer), and the model without squeeze excitation (SE) block (CRMnet without SE block).

Pre-training followed by fine-tuning on the particular dataset demonstrates substantial improvement compared with a model directly trained on the complex medium test set, due to the larger training data set size and improved starting point for fine-tuning model training by transfer learning. Overall performance decreased when the transformer and squeeze excitation blocks were removed, particularly for the random dataset, indicating that both blocks contribute to the model’s predictive performance.

### 3.3 Model interpretation

To explore the biological insights from our trained model, we used saliency maps to interpret the model by visualizing predictive motifs. Saliency maps based on gradient backpropagation have been commonly applied to highlight model-derived features in input data (Adebayo et al., 2018), and have been used to interpret the relationship between the input and prediction of the trained model, where a segment of the input with a higher saliency value indicates an influential region for the model’s prediction (Eraslan et al., 2019). By combining the gradient values with the input sequences, also known as input-masked gradients, we can visualize the segments that significantly impact the model’s prediction (Eraslan et al., 2019).

For comparison, we first searched for significant TF motifs using probabilistic motif discovery based on expression levels (see Methods). We discovered the known yeast motifs associated with higher expression levels: NHP10 (High-mobility group (HMG) domain factors), REB1 (Myb/SANT domain factors), ABF1 (Basic helix-loop-helix factors (bHLH), AZF1 (C2H2 zinc finger factors), and RAP1 (Myb/SANT domain factors) were the top 5 motifs.

We then visualized the input-masked gradients by plotting the saliency map logos generated from our fine-tuned model over yeast native sequences compared to these significant motifs from probabilistic motif discovery. The results show that the saliency map matches the known yeast motif logos (Figure 5 A-E and supplementary figure S2). To quantify this, we further calculated mean saliency map gradients over the positions in the sequences matched by these top 5 motifs and showed that these motifs are associated with substantially higher saliency gradients than the mean over all sequences as a control (Figure 5F).

**Figure 5:**
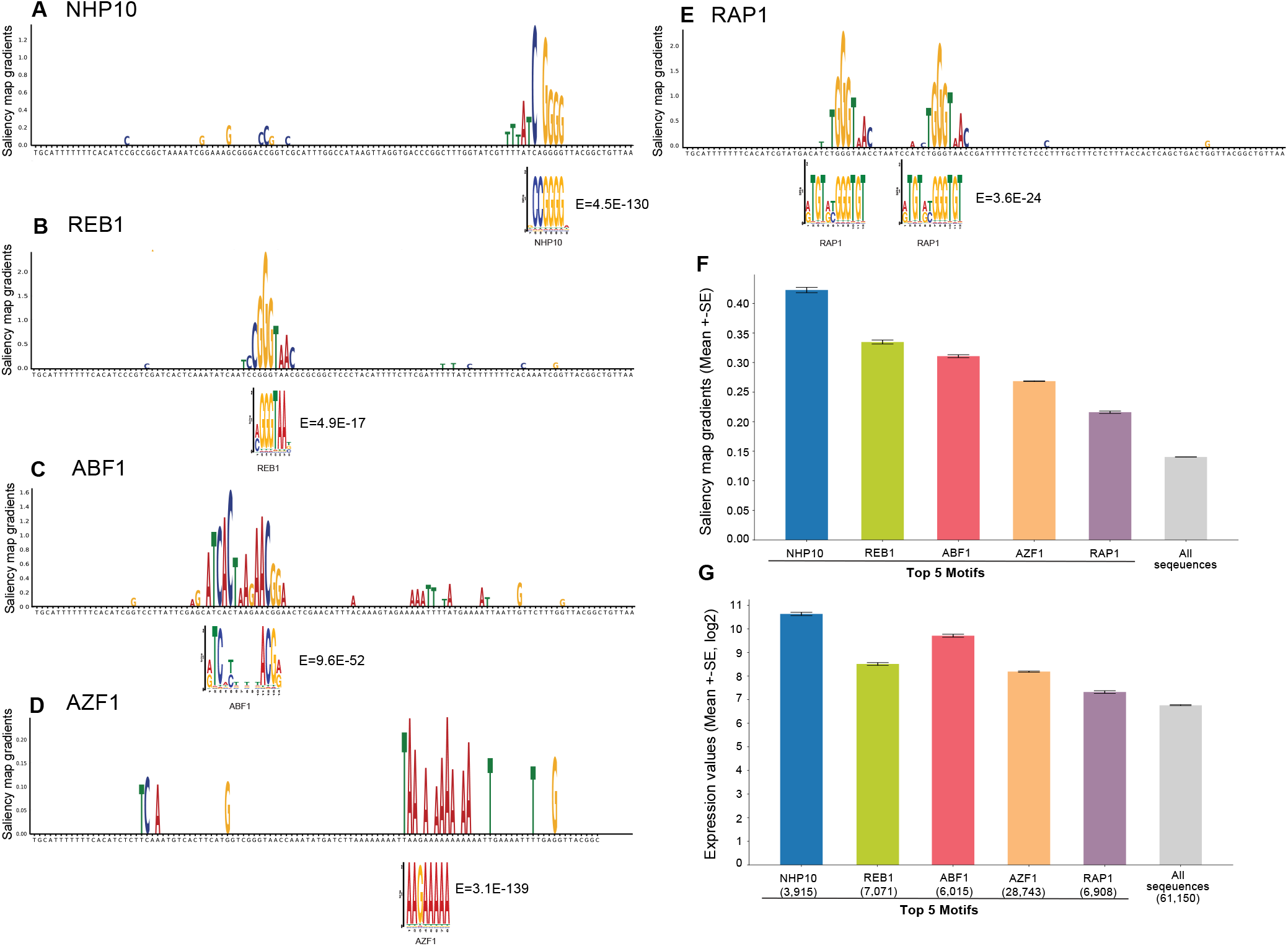
Model interpretation by saliency maps. **A-E:** The top 5 yeast TF motifs detected by motif discovery: NHP10, REB1, ABF1, AZF1, and RBP1. Shown is an example sequence with its saliency map gradients over 80-nt for each motif, aligned with the known TF motif logo and E-values. **F**: Mean saliency map gradients over these top 5 motif matches in yeast native sequences, and mean saliency map gradients over all sequences as the controls. **G**: Mean expression levels of yeast native sequences containing these top 5 TF motifs, and all native sequences as the control.

Furthermore, we calculated the mean expression levels of yeast native sequences containing these top 5 TF motifs compared to all sequences as a control. The result shows that these motifs are associated with higher expression levels as expected (Figure 5G). Notably, the saliency gradients showed that the TF motifs associated with the highest expression levels contribute the most to the prediction of gene expression, supporting that our model extracts biologically meaningful features.

### 3.4 Training time comparisons

We next compared the training time between eight TPU V3 cores and eight GPU A100s under different batch sizes and precision settings (shown in Table 1). Training on the V100 GPU with batch size equal to 1024 and default precision setting was used as the benchmark. The time-per-step is the average processing time to process one batch of data. The average epoch time represents how long it takes to run over all the data. The training time was estimated for the model to run 20 epochs without considering model convergence and the initialization time. The blank value indicates the batch size is too big and over the accelerating hardware’s memory limit.

**Table 1.**
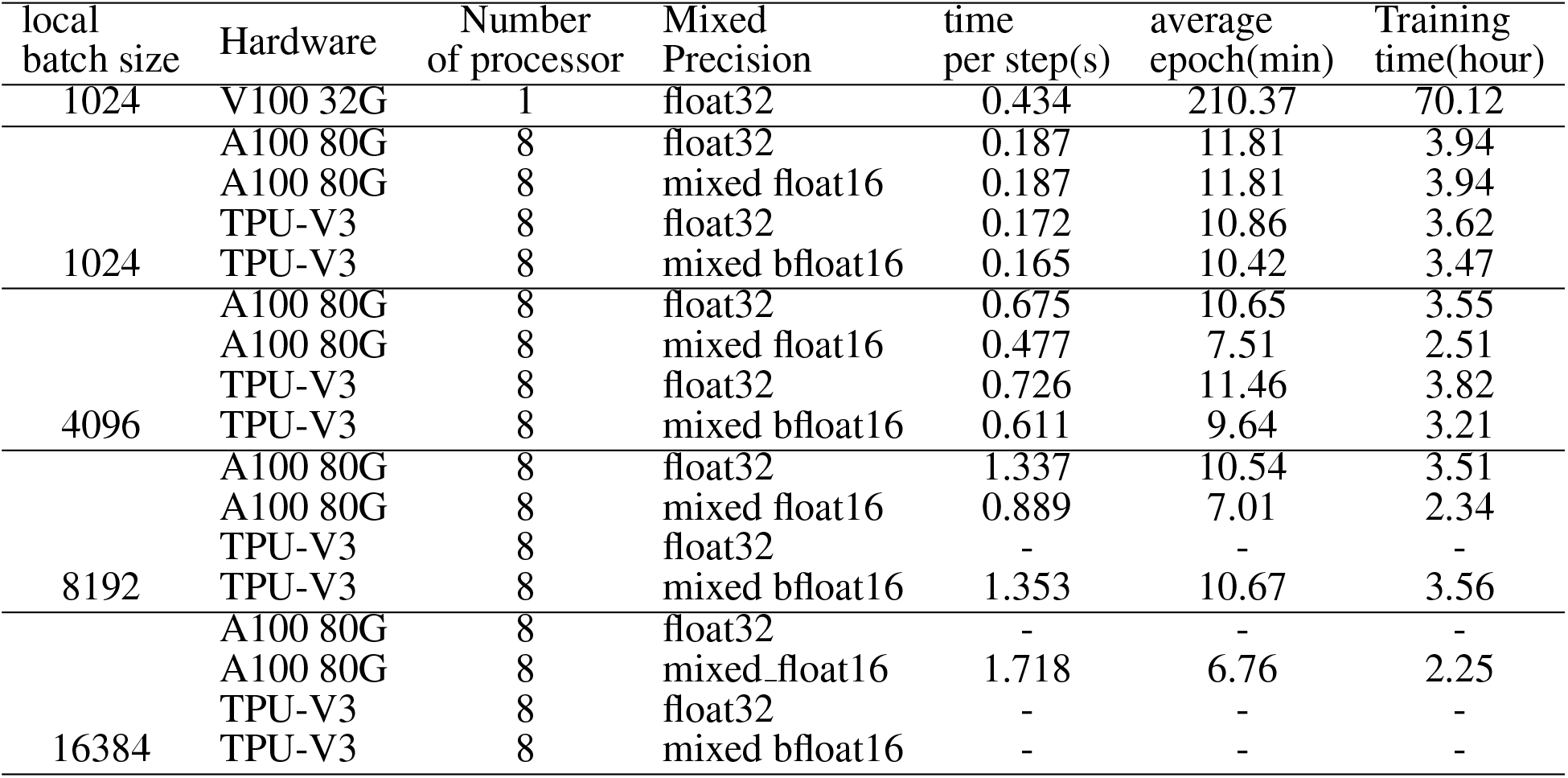
Summary of training speed for different batch sizes and accelerator hardware configurations. The benchmark training time was calculated by training the model on complex medium data with a single V100 GPU, with a local batch size of 1,024 and no mixed precision policy. The final training time is based on training the model for 20 epochs without considering the model’s convergence. The training time was left empty if the local batch was too large and exceeded the hardware’s graphic memory limit. For distributed training on multiple hardware, we compared the training time between TPU v3-8 and DGX box with eight A100s, where each configuration contains eight accelerator hardware.

This study showed that: distributed training on multiple accelerator hardware can reduce training time significantly to feasible levels (from 70h to 4h); mixed precision can improve GPU performance, especially with large batch sizes, and can reduce the memory requirement; and the latest GPU A100 with 80 GB of graphics memory can take input with a larger batch size than TPU v3-8 where each TPU core has 32 GB of memory.

## 4 MATERIALS AND METHODS

### 4.1 Data Collection

In the study of Vaishnav et al. (2022), the yeast cells were grown under different mediums to exercise different metabolic pathways. Here we used data from cells grown in the complex (yeast extract, peptone and dextrose) and defined (lacking uracil) medium as specified in (Vaishnav et al., 2022). The training data was downloaded from https://zenodo.org/record/4436477, containing 30,722,376 and 20,616,659 random sequences from complex and defined medium, respectively, with their expression values evaluated by Gigantic Parallel Reporter Assay experiment (Supplementary table S1).

For model testing data, we used independent test sets drawn from experimental replicate datasets, which were generated in Vaishnav et al. (2022), consisting of native and random promoter sequences (N=61,150 and N=2,954, respectively) from complex medium and (N=3,782 and N=5,284, respectively) from defined medium (Supplementary table S1). The test data was downloaded from GitHub repository (at https://github.com/1edv/evolution).

### 4.2 Model Training Setup

We first trained individual models using data from complex medium and defined medium separately. Specifically, 30,722,376 random sequences from the complex medium and 20,616,659 random sequences from the defined medium were used to train the individual models. For the pre-trained model, we evenly sampled data from both mediums. In total, 51,339,035 sequences and their experimentally measured expression levels were used for pre-training the model. We then trained the pre-train model on complex and defined medium separately in the fine-tuning process.

The model’s performance was assessed using independent test datasets as described above, and none of the sequences in the test datasets were used during model training. It is important to note that the test data library was measured in separate experiments from the training data and that the test data library contains fewer sequences than the experiments used to generate the training data. As a result, the expression value associated with each sequence was precisely measured in the test data (by averaging 100 yeast cells per sequence).

### 4.3 Data Pre-processing

We first used TensorFlow 2 *tf*.*keras*.*preprocessing*.*pad sequence* function to pad the original sequence to a length equal to 112 nt, as the original input sequences do not have a fixed length. We truncated the sequences from the end for sequences longer than 112 nt; for sequences shorter than 112 nt, we padded the sequences from the end with “N”. We then used one-hot encoding to encode the nucleotides based on the order of “A”, “C”, “G”, and “T”. Specifically, we used *tf*.*keras*.*layer*.*StringLookup* function to encode the input sequences and define the vocabulary as [‘A’, ‘C’, ‘G’, ‘T’] while characters not in the vocabulary (i.e., “N”) are encoded in the fifth dimension. Therefore, “A”, “C”, “G”, “T” and “N” are encoded to [1,0,0,0,0], [0,1,0,0,0], [0,0,1,0,0], [0,0,0,1,0] and [0,0,0,0,1], respectively.

### 4.4 Model Execution

Since we mainly used TPU v3-8 VM to train our model, which contains eight tensor processing cores, we set the global batch size to 8192, where each TPU core is assigned 1024 samples. To accelerate the efficiency of feeding data into the model, we prefetch the training data into the memory using the *tf*.*data*.*Dataset*.*prefetch()* function. This operation reduced the latency and improved the data pipeline throughput. And we set the buffer size for prefetching the data equal to *tf*.*data*.*AUTOTUNE*, which will optimize the number of data prefetched automatically.

We distributed the training process of our model in two different hardware: TPUs and GPUs. For Tensor Processing Units (TPU) provided by Google TPU Research Cloud, we trained the model with the TPU v3-8 virtual machine. The TPU v3-8 virtual machine comes with 8 processing cores. So, we set the global batch size equals to 8,192, in which each core is allocated a local minibatch size equal to 1,024. For the GPUs, we trained the model with the Nvidia DGX A100 provided by Australian National Computational Infrastructure, which comes with eight A100 GPUs. Thus, following the same setup with TPUs, we set the global batch size equal to 8,192 to ensure each A100 GPU processes a minibatch with 1,024 samples in parallel.

We used Huber loss to calculate the difference between predictions and true values for the loss function since it is less sensitive to outliers than the mean-square error in regression problems (Huber, 1992). We use Adam optimizer to optimize the Huber loss function. For the learning rate scheduler, we set a learning rate warm-up in the first 10 epochs, which gradually increase the learning rate of the optimizer from 0.0001 x NUM of HARDWARE (i.e., 0.0008) to 0.001 x NUM of HARDWAR (i.e., 0.008). A cosine decay learning rate scheduler was then used to gradually reduce the learning rate 0.0001 x NUM of HARDWARE (i.e., 0.0008). To avoid overfitting, an early stop call-back function was used. This call-back function monitors the model’s performance over the validation dataset. If the model’s validation R-square value is not improved in the most recent ten epochs, it stops training and restores the model weight with the best performance over validation data.

We use the Tensorflow 2 *tf*.*distribute for distributed training*.*MirroredStrategy* to train the model on a DGX A100 box which contains eight A100 GPUs. We used the *tf*.*distribute*.*TPUStrategy* for training on TPU v3-8 virtual machine, which has eight tensor cores. Both strategies are synchronous training processes intended to distribute training across multiple processing units on a single machine. The synchronous strategy first copied all of the model’s variables to each processor. The gradients from each processor were then fused using all-reduce. The resulting value will be synchronized to all instances stored in each processor. Since our model training does not require high precision and training on TPUs automatically uses float16, so for training on GPUs, we use the mix precision policy by setting up precision equal to mixed float16. The mixed precision policy improves our model training speed on Amber GPUs without losing accuracy.

We further compared the training speed between A100 GPUs and TPU V3-8 with different local batch sizes and mixed precision policy. To reduce the impact of other factors, such as the time used to build the computational graph, we exclude the time reported in the first epoch and take the maximum value from the time-per-step column. The time-per-step report is the average time the hardware processes each batch in one epoch. The training speed comparison is shown in Figure 1. Note that the final training time is an optimistic estimation for training the model for 20 epochs, which doesn’t guarantee the model coverage and doesn’t consider the overhead time used for inter-core communication.

For the software and packages, we used Python 3.8.10 to write the code for training and evaluation. For data pre-processing, we mainly use NumPy 1.22.1 and Pandas 1.5.0 packages. For building a deep learning model, we use TensorFlow 2.8.0 framework to implement and train the neural network. Functions from TensorFlow Addons 0.16.1 and SciPy 1.9.3 are used to evaluate models’ performance. Matplotlib 3.6.1 and Seaborn 0.12.1 are used for visualization.

### 4.5 Motif Discovery

We used the motif discovery tool MEME suite (Bailey et al., 2015) “Differential Enrichment mode” to detect the motif enrichment in the top 2000 sequences with high gene expression against the bottom 2000 sequences with low gene expression. We used FIMO in the MEME suite to search for motif hits in yeast native promoter sequences.

We further used two ranking-based methods, Discovering Ranked Imbalanced Motifs using Suffix Trees (DRIMust) (Leibovich et al., 2013) and rGADEMm (Mercier et al., 2011), to determine the*de novo* motifs in a ranked list of sequences, which were ranked from high to low expression values. DRIMust uses suffix trees to identify overrepresented motifs in the top-ranked sequences and further evaluates the obtained k-mers by minimum-hypergeometric (mHG) approach (Leibovich et al., 2013). rGADEM combines spaced dyads and an expectation-maximization (EM) algorithm. The spaced dyads are identified by their overrepresentation in the input sequences, and a genetic algorithm is further employed to mark them significant and to declare them as motifs (Mercier et al., 2011). We used Bioconductor packages “TFBStools” (Tan and Lenhard, 2016), “JASPAR” (Castro-Mondragon et al., 2022), to match the identified motifs with known JASPAR Yeast motifs.

We selected the top 5 motifs ranked by E-value in MEME that were reproduced by the rank-based methods.

## 5 CONCLUSION

In this study, we introduced CRMnet, a novel neural network architecture that accurately predicts the gene expression levels of yeast promoter sequences. First, we adopted the U-Net architecture from the image semantic segmentation task and applied it to genomic sequences as a feature extractor. Furthermore, we utilized transformer encoders, which leverage self-attention mechanisms to extract additional useful information from genomic sequences. The extracted features were then fed into an MLP to predict the expression levels as a regression problem. By testing on data not used during the training process, our model surpassed the benchmark networks of Vaishnav et al. (2022). Our ablation studies of the CRMnet model demonstrated the potential for improvements in predictive performance for a given biological problem by the design of custom DNN architectures. In particular, augmentation of a model with a combination of CNN and additional transformer stages guided by training and testing results on large high-throughput datasets can give useful increments in performance.

Importantly, high performance DNN models extracting dependency information via attention mechanisms allow for biological insights through model interpretation. In this study, we visualized regions of key importance for expression regulation by plotting the saliency map over the input yeast DNA sequences. Notably, we found that the logo plots constructed from saliency maps over the input sequences are correlated with the sequence motifs of known yeast transcription factors. Future improvements in DNN model architectures along with improved model interpretation methods will be key enablers for future biological discoveries of subtle regulatory signals.

## 7 DATA AVAILABILITY STATEMENT

The code is available on GitHub at https://github.com/jiayuwen/CRMnet. The final model and prepossessed data are available at https://zenodo.org/record/7375243#.Y4gDjS0RoUE.

## 8 ACKNOWLEDGMENTS

This study was supported by the National Computational Infrastructure (NCI Australia) and Google’s TPU Research Cloud (TRC). We thank Dr. Jingbo Wang from NCI for the support.

## 10 AUTHOR CONTRIBUTIONS

KD conducted all machine learning coding and drafted the manuscript. GD collected the data and performed the motif analysis. BJP and JW supervised the project, provided guidance, discussed results and revised the manuscript. All authors read the current manuscript and approved the submitted version.

## 12 SUPPLEMENTAL DATA

The Supplementary Material for this article can be found at: “supplementary CRMnet.pdf”

## REFERENCES

Adebayo, J., Gilmer, J., Muelly, M., Goodfellow, I., Hardt, M., and Kim, B. (2018). Sanity checks for saliency maps. Advances in neural information processing systems 31

Bailey, T. L., Johnson, J., Grant, C. E., and Noble, W. S. (2015). The meme suite. Nucleic acids research 43, W39–W49

Castro-Mondragon, J. A., Riudavets-Puig, R., Rauluseviciute, I., Berhanu Lemma, R., Turchi, L., Blanc-Mathieu, R., et al. (2022). Jaspar 2022: the 9th release of the open-access database of transcription factor binding profiles. Nucleic acids research 50, D165–D173

Chen, J., Lu, Y., Yu, Q., Luo, X., Adeli, E., Wang, Y., et al. (2021). Transunet: Transformers make strong encoders for medical image segmentation. arXiv preprint arXiv:2102.04306

Davidson, E. H. and Erwin, D. H. (2006). Gene regulatory networks and the evolution of animal body plans. Science 311, 796–800

de Boer, C. G., Vaishnav, E. D., Sadeh, R., Abeyta, E. L., Friedman, N., and Regev, A. (2020). Deciphering eukaryotic gene-regulatory logic with 100 million random promoters. Nature biotechnology 38, 56–65

Dosovitskiy, A., Beyer, L., Kolesnikov, A., Weissenborn, D., Zhai, X., Unterthiner, T., et al. (2020). An image is worth 16×16 words: Transformers for image recognition at scale. arXiv preprint arXiv:2010.11929

Eraslan, G., Avsec, Ž., Gagneur, J., and Theis, F. J. (2019). Deep learning: new computational modelling techniques for genomics. Nature Reviews Genetics 20, 389–403

Hu, J., Shen, L., and Sun, G. (2018). Squeeze-and-excitation networks. In Proceedings of the IEEE conference on computer vision and pattern recognition. 7132–7141

Huber, P. J. (1992). Robust estimation of a location parameter. In Breakthroughs in statistics (Springer). 492–518

Leibovich, L., Paz, I., Yakhini, Z., and Mandel-Gutfreund, Y. (2013). Drimust: a web server for discovering rank imbalanced motifs using suffix trees. Nucleic acids research 41, W174–W179

Li, J., Pu, Y., Tang, J., Zou, Q., and Guo, F. (2021). Deepatt: a hybrid category attention neural network for identifying functional effects of dna sequences. Briefings in bioinformatics 22

Mathelier, A., Shi, W., and Wasserman, W. W. (2015). Identification of altered cis-regulatory elements in human disease. Trends in Genetics 31, 67–76

Mercier, E., Droit, A., Li, L., Robertson, G., Zhang, X., and Gottardo, R. (2011). An integrated pipeline for the genome-wide analysis of transcription factor binding sites from chip-seq. PloS one 6

Ni, P. and Su, Z. (2021). Accurate prediction of cis-regulatory modules reveals a prevalent regulatory genome of humans. NAR genomics and bioinformatics 3

Quang, D. and Xie, X. (2016). Danq: a hybrid convolutional and recurrent deep neural network for quantifying the function of dna sequences. Nucleic acids research 44, e107–e107

Ronneberger, O., Fischer, P., and Brox, T. (2015). U-net: Convolutional networks for biomedical image segmentation. In International Conference on Medical image computing and computer-assisted intervention (Springer), 234–241

Springenberg, J. T., Dosovitskiy, A., Brox, T., and Riedmiller, M. (2014). Striving for simplicity: The all convolutional net. arXiv preprint arXiv:1412.6806

Stewart, A. J., Hannenhalli, S., and Plotkin, J. B. (2012). Why transcription factor binding sites are ten nucleotides long. Genetics 192, 973–985

Tan, G. and Lenhard, B. (2016). Tfbstools: an r/bioconductor package for transcription factor binding site analysis. Bioinformatics 32, 1555–1556

Vaishnav, E. D., de Boer, C. G., Molinet, J., Yassour, M., Fan, L., Adiconis, X., et al. (2022). The evolution, evolvability and engineering of gene regulatory dna. Nature 603, 455–463

Vaswani, A., Shazeer, N., Parmar, N., Uszkoreit, J., Jones, L., Gomez, A. N., et al. (2017). Attention is all you need. Advances in neural information processing systems 30

Wang, Y., Wei, G.-Y., and Brooks, D. (2020). A systematic methodology for analysis of deep learning hardware and software platforms. Proceedings of Machine Learning and Systems 2, 30–43

You, K., Liu, Y., Wang, J., and Long, M. (2021). Logme: Practical assessment of pre-trained models for transfer learning. In International Conference on Machine Learning (PMLR), 12133–12143

Zhou, J. and Troyanskaya, O. G. (2015). Predicting effects of noncoding variants with deep learning–based sequence model. Nature methods 12, 931–934

