## Supplemental_Materials for "CRMnet: a deep learning model for predicting gene expression from large regulatory sequence datasets"

### *Supplementary Material*

#### 1 SUPPLEMENTARY TABLES AND FIGURES

##### 1.1 Tables

**Table S1.** Summary of yeast training/testing datasets.

| Data type | Usage | Yeast medium | #sequences |
|---|---|---|---|
| Random Promoter Sequences | training/testing | Complex (YPD) | 30,722,376 |
| Random Promoter Sequences | training/testing | Defined (SD-Uracil) | 20,616,659 |
| Native Yeast Promoter Sequences | testing | Complex (YPD) | 61,150 |
| Random Promoter Sequences | testing | Complex (YPD) | 2,954 |
| Native Yeast Promoter Sequence | testing | Defined (SD-Uracil) | 3,782 |
| Random Promoter Sequences | testing | Defined (SD-Uracil) | 5,289 |

##### 1.2 Figures

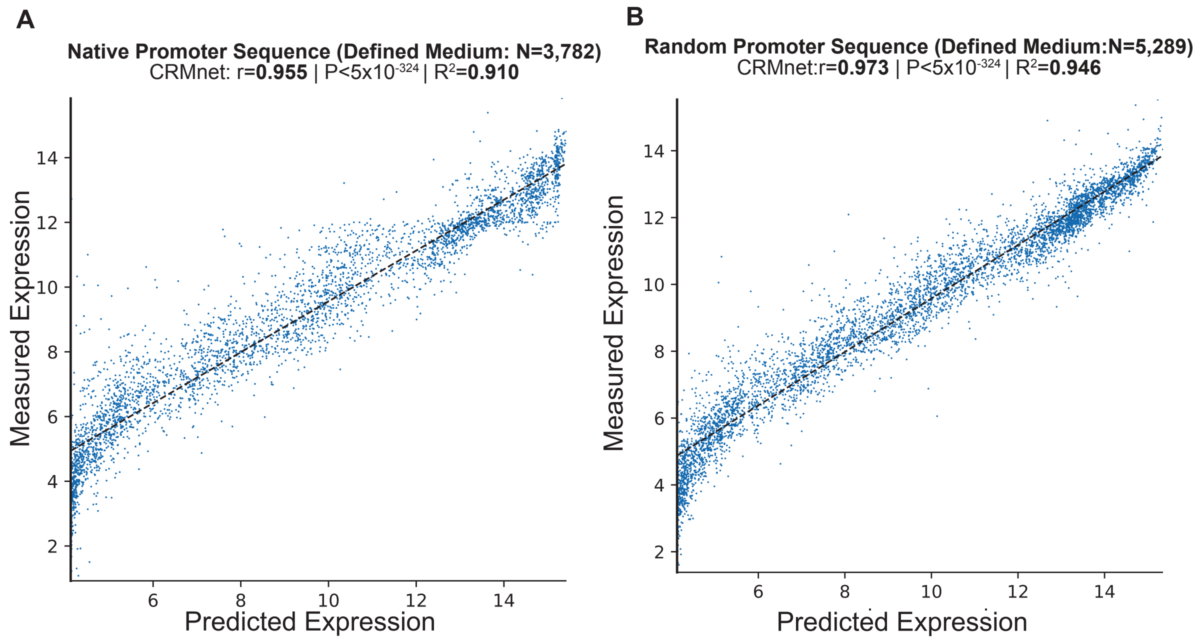


**Figure S1. Prediction of expression from yeast native sequences in defined medium from the fine-tune CRMnet.** Fine-tuned CRMnet tested on A: native promoter sequences; and B. random promoter sequences. The y-axes represent measured expression levels, while the x-axes represent predicted expression levels. As a benchmark, the model performance metrics of the Pearson $r$ value, associated two-tailed p-values, and R-square for the transformer model from Vaishnav et al. (2022) showed: A: r=0.950, P $< 5 \times 10^{-324}$, $R^2$=0.900; B: r=0.968, P $< 5 \times 10^{-324}$, $R^2$=0.937

#### REFERENCES

Vaishnav, E. D., de Boer, C. G., Molinet, J., Yassour, M., Fan, L., Adiconis, X., et al. (2022). The evolution, evolvability and engineering of gene regulatory dna. *Nature* 603, 455–463

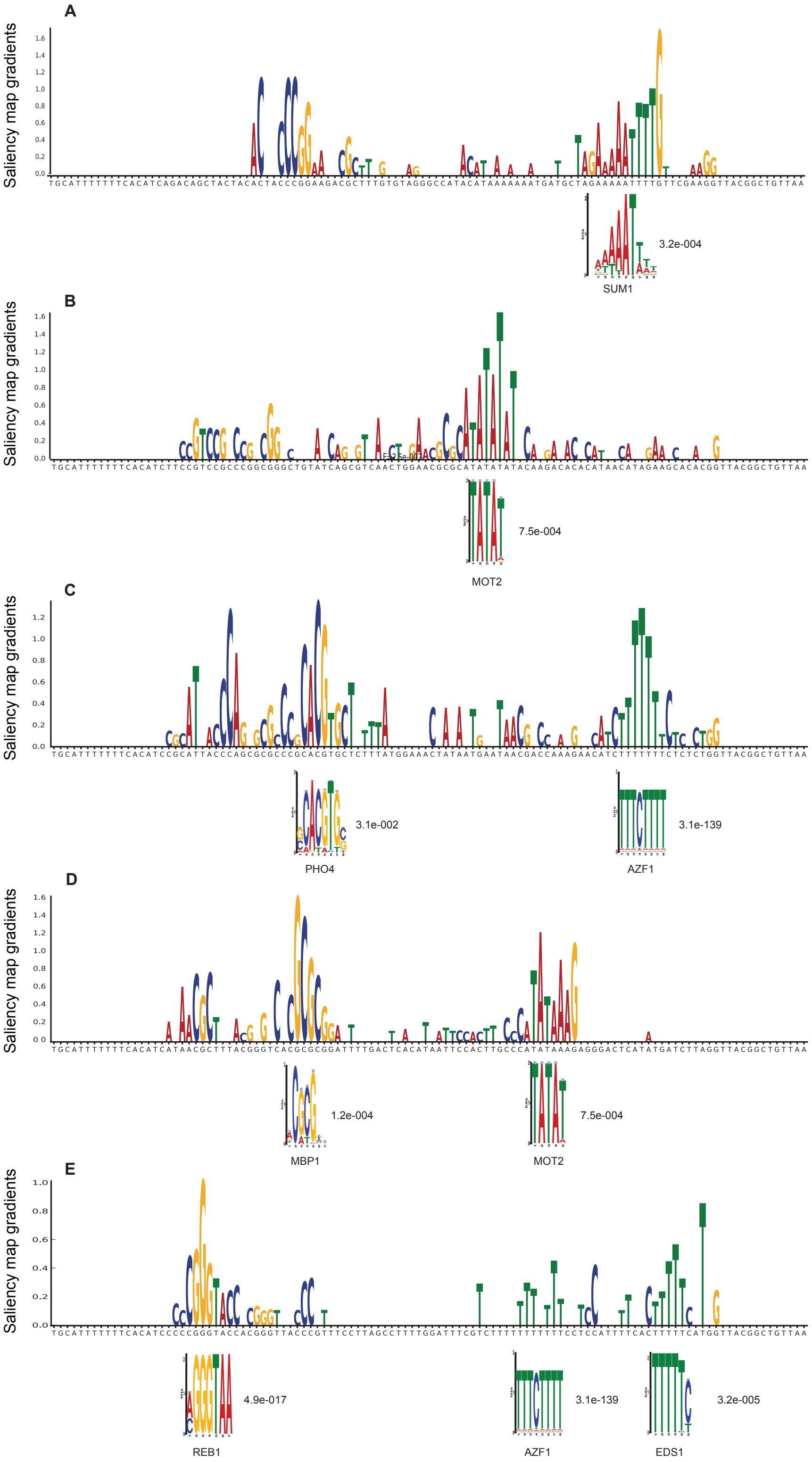


**Figure S2. Model interpretation by saliency maps.** Additional TF motifs detected by motif discovery (E-values < 1E-3). Shown is an example sequence with its saliency map gradients over 80-nt for each motif, aligned with the known TF motif logo and E-values.
